# Detection of four imperiled western North American freshwater mussel species from environmental DNA with multiplex qPCR assays

**DOI:** 10.1101/2020.03.27.012088

**Authors:** Torrey W. Rodgers, Joseph C. Dysthe, Cynthia Tait, Thomas W. Franklin, Michael K. Schwartz, Karen E. Mock

## Abstract

We developed multiplexed, species-specific, quantitative PCR assays for the detection of four freshwater mussel species native to western North America, *Gonidea angulata, Margaritifera falcata*, *Anodonta nuttalliana* and *Anodonta oregonensis*, from environmental DNA (eDNA). These species have experienced dramatic declines over the last century and are currently threatened in many portions of their ranges. Therefore, improved tools for detecting and monitoring these species are needed. Species-specificity and sensitivity of assays were empirically tested in the lab, and multiplex assays were also validated with field collected eDNA samples. All assays were species-specific, sensitive, and effective for detection from eDNA samples collected from streams and rivers. These assays will aid in the detection, monitoring, management, and conservation of these vulnerable species.

## Introduction

Western North America is home to five native freshwater mussel species, *Gonidea angulata, Margaritifera falcata*, *Anodonta oregonensis, Anodonta nuttalliana, and Sinanodonta beringiana*. Originally, as many as six separate *Anodonta* species were described from western North America based on morphology, but genetic analyses have shown that only three major genetic lineages exist (Chong et al. 2008). *Sinanodonta beringiana*, which was recently reassigned from *Anodonta* to *Sinanodonta* (Williams et al. 2017), is only known from Alaska and Canada, although the southern extent of its range is uncertain. The remaining two *Anodonta* lineages in North America include a clade comprised of *Anodonta oregonensis* (I. Lea, 1838) and *Anodonta kennerlyi* (I. Lea, 1860) (here referred to as *A. oregonensis*), and a clade comprised of *Anodonta nuttalliana* (I. Lea, 1838), *Anodonta californiensis* (I. Lea, 1852), *and Anodonta wahlamatensis* (I. Lea 1938) (here referred to as *A. nuttalliana*). *A. oregonensis* ranges from Alaska to northern California (Blevins et al. 2017), and *A. nuttalliana* ranges from Washington State to northern Mexico, extending east into Idaho, Wyoming, Utah, Nevada and Arizona (Blevins et al. 2016a). *Gonidea angulata* occurs in California, Nevada, Oregon, Washington, Idaho, and British Columbia, while *M. falcata* (Gould, 1850) inhabits these states plus parts of Wyoming, Montana, Utah, and Alaska (Blevins et al. 2017).

Native freshwater mussel species have experienced dramatic declines in both distribution and abundance in Western North America, due primarily to human impacts such as habitat degradation and introduction of non-native species (Haag 2012, Strayer 2014). As filter feeders, these species provide important ecosystem services such as improving water clarity, and are a food source for wildlife species. Ongoing population declines highlight the importance of monitoring native freshwater mussels, but traditional sampling requires time-consuming surveys, often necessitating snorkeling or SCUBA diving. Additionally, species identification requires trained expertise, and for *Anodonta* species can even necessitate genetic analyses. Thus, improved tools for monitoring these species more efficiently and economically are needed.

Environmental DNA (eDNA) has emerged as an efficient and reliable tool for aquatic species monitoring, has been shown to be more sensitive than traditional methods under most conditions for fish (Wilcox et al. 2016), amphibians (Smart et al. 2015), reptiles (Hunter et al. 2015), and has recently been used to detect freshwater mussels (Stoeckle et al. 2016, Currier et al. 2018, Dysthe et al. 2018). We designed new, species-specific eDNA assays for *A. nuttalliana*, *A. oregonensis*, and *G. angulata*, and developed three multiplexed quantitative PCR (qPCR) assays that employ these along with an assay previously designed for *M. falcata* (Dysthe et al. 2018). The first multiplex assay detects and discerns *A. nuttalliana* and *A. oregonensis*, the second multiplex assay detects and discerns *G. angulata* and *M. falcata*, and the third multiplex assay detects and discerns *A. nuttalliana* and *M. falcata.* We also tested the ability of all three multiplex assays for species detection from field collected, filtered water samples. We did not design an assay for *S. beringiana* because this species does not currently appear to face the same conservation challenges as the other four North American species (Vinarski and Cordeiro 2011), and because available tissues and sequence data are currently insufficient for design of a robust assay for this species.

## Methods

### Assay Design

For assay design, sequence data for *A. nuttalliana, A. oregonensis, and G. angulata* spanning the species’ ranges from the mitochondrial gene cytochrome oxidase subunit 1 (COI) were compiled from our in-house freshwater mussel sequence database (sequences from 344 samples from 67 populations for *A. nuttalliana*, 95 samples from 23 populations for *A. oregonensis*, and 145 samples from 31 populations for *G. angulata;* Table S1). Additional sequences from GenBank were also included, (27 sequences for *A. nuttalliana*, 11 sequences for *A. oregonensis*, and 6 sequences for *G. angulata;* Table S2; accessed Nov 2018). For *A. nuttalliana* we included sequences from Washington (WA; n=25), Oregon (OR; n=112), Idaho (ID; n=12), Utah (UT; n=32), Nevada (NV; n=20), California (CA; n=147), Arizona (AZ; n=9) and Mexico (n=11). For *A. oregonensis* we included sequences from OR (n=51), WA (n=31), CA (n=6), British Columbia (BC; n=11) and Alaska (AL; n=3; Figure 1, Tables S1 and S2). For *G. angulata*, we included sequences from WA (n=24) OR (n=36), NV (n=10), CA (n=40), BC (n=26) and ID (n=12). COI sequence data from the non-target native species *S. beringiana* were obtained from our in-house sequence database, and COI sequence data from five other potentially sympatric, non-target, taxa: *Corbicula fluminea*, *Dreissena polymorpha, Dreissena bugensis, Ferrissia rivularis*, and *Lampsilis siliquoidea*, were retrieved from GenBank (Table S3).

**Figure 1.**
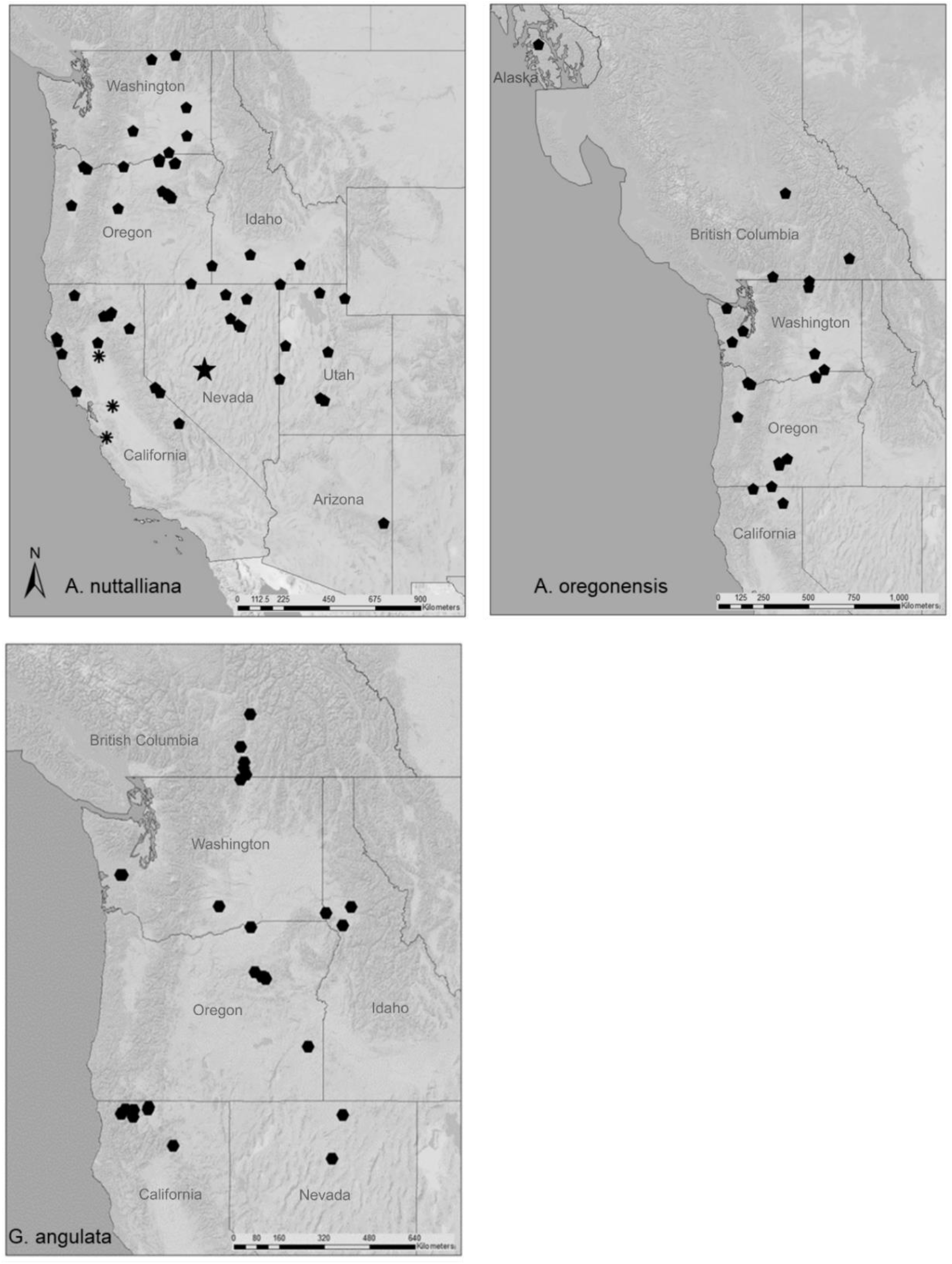
Locations in western North America of sample sequences used to design species-specific qPCR assays for the freshwater mussel species *Anodonta nuttalliana, Anodonta oregonensis*, and *Gonidea angulata.* Asterisk designates locations were sample sequences match probe AnuCOI2-Pc. Star designates location where 1 sample sequence did not match probe AnuCOI2-Pb or AnuCOI2-Pc. At all remaining locations sequences were a perfect match to the preferred assay for each species.

Sequence data from all species were aligned using Sequencher software v5.2 (Gene Codes, Ann Arbor, MI), and species-specific primers were generated with the online tool DECEPHER (Wright et al. 2014). We then used ABI primer express software v3.0.1 (Applied Biosystems, Foster City, CA) to design TaqMan™ Minor-Groove-Binding (MGB) qPCR probes. For *A. nuttalliana*, we initially tested two candidate primer sets and three TaqMan™ probes (one probe for primer set one, and two for primer set two; Table 1). We later tested an additional probe for use in the Central Valley of CA where a SNP (single nucleotide polymorphism) in the probe region was identified for some populations (see results). For *A. oregonensis*, we tested one candidate primer set and two TaqMan™ probes. For *G. angulata* we initially tested 3 candidate primer sets in SYBR^®^ green qPCR, and then one TaqMan™ probe (Table 1). For *M. falcata*, our multiplex assays utilized an existing single-species assay described in detail in Dysthe et al. (2018).

**Table 1.**
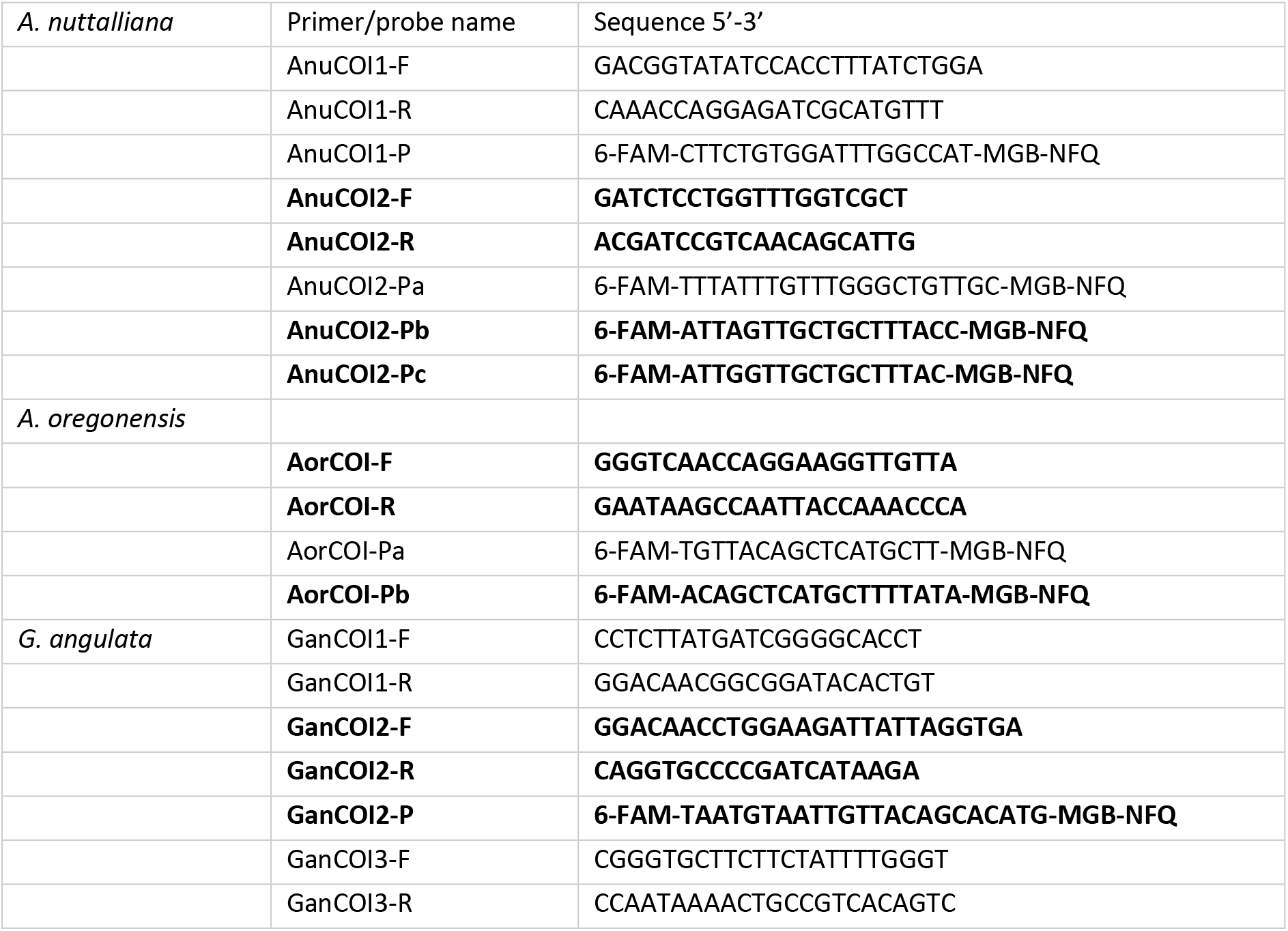
Primer and probe sets developed and initially tested for detection of *A. nuttalliana, A. oregonensis*, and *G. angulata*. Primers and probes in bold were chosen as preferable for use in most populations after testing and were used in multiplex assays.

### Specificity testing

Specificity testing of our assays included both *in silico* and *in vitro* components. To evaluate primer specificity *in silico*, we used NCBI Primer-BLAST (Ye et al. 2012) against the NCBI nr database to reduce likelihood that species occurring in western North America could potentially cross-amplify and produce false positives.

In the laboratory, we initially tested specificity of candidate primer sets using SYBR^®^ green qPCR. Each primer set was tested with tissue DNA extracts from each target species, as well as four DNA extracts from each non-target species (S*. beringiana, A. nuttalliana, A. oregonensis*, *G. angulata* and *M. falcata*, depending on the assay being tested). qPCR reactions included 7.5 μl Power SYBR^®^-Green Mastermix (Thermo-Fisher Scientific), 900 nM of each primer, and 0.1 ng of template DNA in a total reaction volume of 15 μl. Cycling conditions were 95ᵒ for 10 minutes followed by 45 cycles of 95ᵒ for 15 seconds and 60ᵒ for one minute, followed by a melt curve.

We then tested primer sets and MGB probes in TaqMan™ qPCR. As above, analyses included five DNA extracts from each target species, and four DNA extracts from each non-target species (S*. beringiana, A. nuttalliana, A. oregonensis*, *G. angulata* and *M. falcata*, depending on the assay being tested). qPCR reactions included 7.5 μl TaqMan™ Environmental Mastermix 2.0 (Thermo-Fisher Scientific) 900 nM of each primer, 250 nM of probe, and 0.1 ng of template DNA in a total reaction volume of 15 μl. Cycling conditions were 95ᵒ for 10 minutes followed by 45 cycles of 95ᵒ for 15 seconds and 60ᵒ for one minute. For further optimization and sensitivity testing, we selected a single primer set and probe for each species based on performance (see results). Selected assays included primer set AnuCOI2 and probe AnuCOI2-Pb for *A. nuttalliana*, primer set AorCOI and probe AorCOI-Pb for *A. oregonensis*, and primer set GanCOI2, and probe GanCOI2-P for *G. angulata* (Table 2). Specificity testing for all assays was conducted in singleplex with 6-FAM-labelled probes (Table 1), prior to testing sensitivity and field validation with multiplex assays including 6-FAM- and NED-labelled probes (Table 2).

**Table 2.**
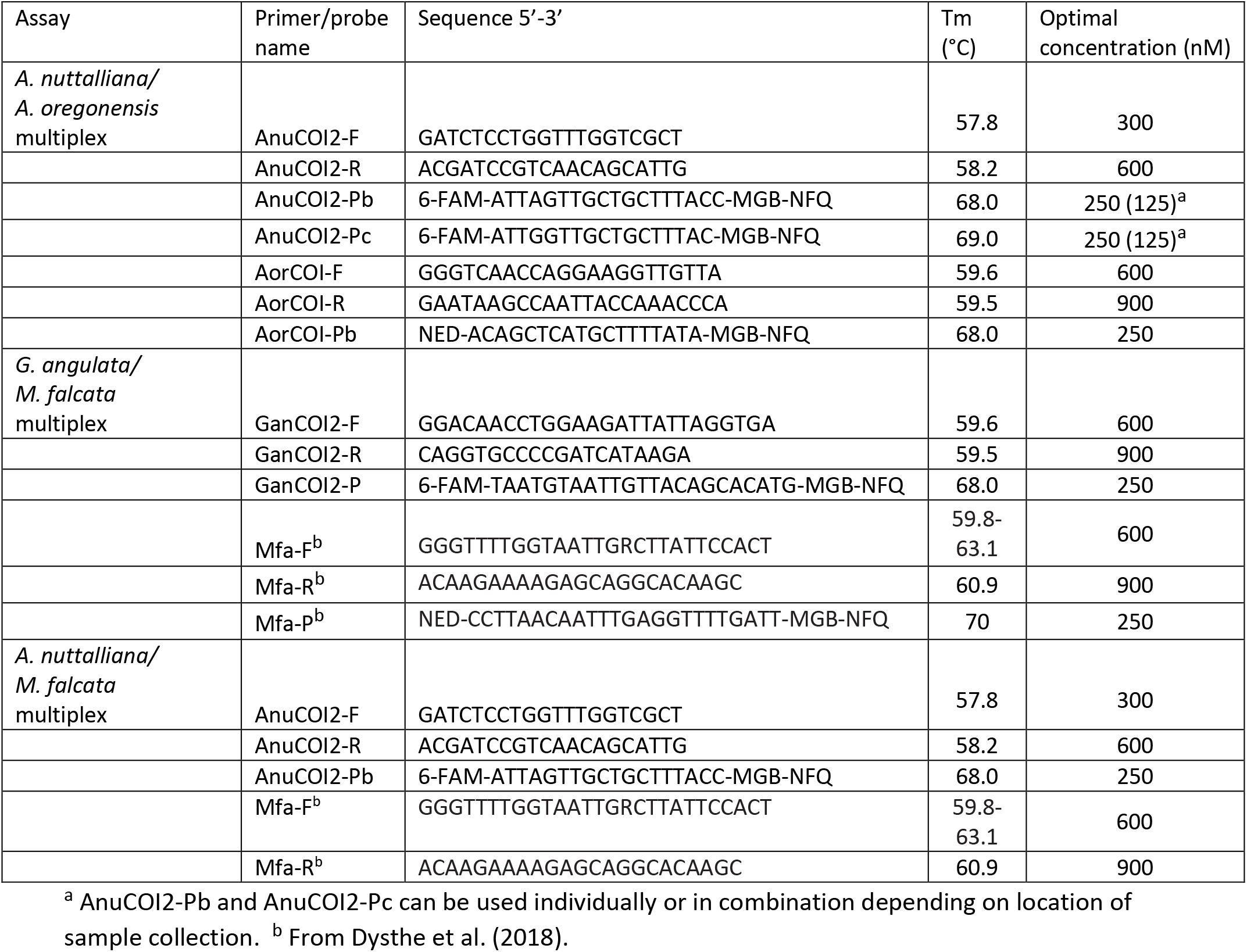
Primers and probes used in multiplex assays for detection of *Anodonta nuttalliana, A. oregonensis, Gonidea angulata*, and *Margaritifera falcata* from environmental DNA.

We also tested an additional TaqMan™ probe (Probe AnuCOI2-Pc; Table 2) for *A. nuttalliana* for use in central CA populations which possessed a SNP in the initial probe (see results). We tested this modified probe on tissue samples from the Pajaro (n=1), Sacramento (n=1), and San Joaquin (n=4) rivers (sites ASA, ASJ, ASM, ASV, ASQ, and APJ; Table S1). In addition, we tested the above tissue samples, as well as tissue samples from ID, OR and UT with both probes AnuCOI2-Pb and AnuCOI2-Pc combined at 125 nM each.

### Primer concentration optimization

Primer concentrations were optimized by running varying concentrations of each forward and reverse primer (100 nM, 300 nM, 600 nM, and 900 nM, each in triplicate), with 100 copies of target-species MiniGene synthetic plasmid DNA (Integrated DNA Technologies, Coralville, IA) per reaction. Primer concentrations with the greatest peak fluorescence and lowest Ct value were selected for subsequent analyses (Table 2).

### Sensitivity testing

To test assay sensitivity, we used MiniGene synthetic plasmids containing the assay sequence for each assay. Plasmids were suspended in 100 μL of IDTE (10 mM Tris, 0.1 mM EDTA) buffer, linearized by digestion with the enzyme Pvu1, and then purified with a PureLink PCR Micro Kit (Invitrogen, Carlsbad, CA) following standard protocols. The resulting products were quantified on a Qubit fluorometer, and estimated concentrations were converted to copy number based on molecular weight (Wilcox et al. 2013). These products were then diluted in IDTE buffer to create quantities of 2, 5, 10, 20, 50, and 100 copies/reaction, and each quantity was analyzed in ten qPCR replicates for each species to determine assay sensitivity for each multiplex.

### Field validation

We tested all three multiplex assays *in vivo* using eDNA samples collected from water bodies where one or more of our target species was historically present. Traditional sampling was not conducted at most sites, but our dataset included a subset of positive control sites where live mussels were documented either at the time of, or within one year of, eDNA sampling (16 sites for *Anodonta*, 15 sites for *M. falcata*, and 5 sites for *G. angulata*; Tables S4 and S5). For the *Anodonta* multiplex, we analyzed 70 eDNA samples from 36 water bodies from 7 states (Table S4). For the *M. falcata/G. angulata* multiplex, we analyzed 31 eDNA samples from 24 water bodies from 6 states (Table S5). For the *A. nuttalliana/M. falcata* multiplex we analyzed 29 eDNA samples from 22 water bodies from 6 states (all of which were positive for either *A. nuttalliana*, *M. falcata*, or both, with the other multiplex assays; Table S6).

Environmental DNA samples were either collected specifically for this project, or they were repurposed from other previously collected samples. One benefit of eDNA sampling is that a sample can be re‐analyzed to detect additional taxa without additional field effort even when initially collected to detect a different species (Franklin et al. 2019, Dysthe et al. 2018). All samples were collected following the protocol outlined in Carim et al. (2016a). Up to five liters of water were pumped through a 1.5 μm glass microfiber filter using a peristaltic pump, and filters were stored in silica desiccant until laboratory processing. We additionally collected and analyzed ‘field blank’ filter samples (n=7) consisting of one liter of distilled water brought from the lab and filtered in the field to act as a negative control to monitor for field-based cross contamination. Samples were extracted in a room dedicated for this purpose using the DNeasy Blood & Tissue Kit (Qiagen, Inc. Valencia, CA, USA) with a modified protocol described in Carim et al. (2016b). Each round of eDNA extraction (n=8) included one ‘extraction blank’ negative control consisting of a clean filter. All samples were analyzed in triplicate qPCR with 4 μl of eDNA extract, 7.5 μl TaqMan™ Environmental Mastermix 2.0 (Thermo-Fisher Scientific), optimized primer and probe concentrations (Table 2), and cycling conditions described above. Each qPCR run included three to six ‘no template’ negative control reactions to monitor for cross-contamination. eDNA extractions were conducted in a dedicated clean lab space physically separated from post-PCR spaces and tissue DNA extraction spaces, and all qPCR reactions were set up under a dedicated PCR hood sterilized with UV radiation prior to each qPCR run (Goldberg et al. 2016).

## Results

### Assay design

The species-specific assays we developed target a 125 bp, 128 bp, and 148 bp fragment of COI in *A. nuttalliana, A. oregonensis*, and *G. angulata* respectively. For *A. oregonensis* and *G. angulata*, the assays were a perfect match at both primers and the probe for all haplotypes from our database and GenBank across the species’ ranges. For *A. nuttalliana*, we were not able to design a single assay that perfectly matched all haplotypes across the species range due to high levels of haplotype diversity. However, we were able to design one assay that perfectly matched all sequences in our database from AZ, ID, OR, UT, and WA. One *A. nuttalliana* individual from one site in NV (Reese River; site ARR from supplementary Table S1) displayed a single SNP in probe AnuCOI2-Pb, but this sequence was a perfect match to probe AnuCOI2-Pa. Thus, use of probe AnuCOI2-Pa may be preferable in this population. A second sample from the Reese River, and all other haplotypes from NV, were a perfect match to both probes. In CA, two *A. nuttalliana* samples possessed single SNPs in the reverse primer. A single sample from the San Joaquin River (Site ASV; Table S1) possessed one SNP in the reverse primer six bp from the 3’ end, but seven other samples collected at the same location were a perfect match to the primer. Likewise, a single sample from the Owens River (Site AOR; Table S1) also possessed one SNP in the reverse primer 14 bp from the 3’ end, but nine other samples from the same location were a perfect match to the primer. Such mismatches far from the 3’ end typically have minimal effect on assay sensitivity (Lefever et al. 2013). Of the 11 *A. nuttalliana* reference sequences on GenBank from Mexico (Table S2), five were a perfect match to the assay, whereas the remaining six possess one SNP 13 bp from the 3’ end of the forward primer and a second SNP 8 bp from the 3’ end in the reverse primer. Thus, if eDNA work is to be conducted in Mexico, a modification of the assay may be warranted.

Additionally, all *A. nuttalliana* samples from the Sacramento, San Joaquin, and Pajaro rivers in central CA (Sites ASA, ASJ, ASM, ASV, ASQ, and APJ; Table S1 and Figure 1) possessed a single guanine-adenine (G-A) SNP in the probe for AnuCOI2-Pb, 15 bp from the 3’ end. This SNP appears to be monomorphic in these populations: all 32 samples from these rivers possessed this SNP. Thus, we designed an additional modified probe with a G instead of an A in the position of the SNP (probe AnuCOI2-Pc).

### Specificity testing

*In silico* evaluation of primer specificity with NCBI Primer-BLAST for *A. nuttalliana* identified just one species with the potential to cross-amplify with our chosen primer set, the congener *A. impura*, which is native to central Mexico and is not known to occur in western North America (MolluscaBase 2019). For *A. oregonensis*, Primer-BLAST identified 14 mussel species with the potential to cross-amplify (Table S7), but none of these taxa occur in western North America. For *G. angulata*, no species with a potential to cross-amplify were identified.

The *in vitro* evaluation of primer specificity using SYBR^®^ Green indicated primer sets for all three species were species-specific, amplifying DNA from all targets at least 10 cycles earlier than DNA from any non-target species, a range which is suitable for specificity once a TaqMan™ MGB probe is added. For *A. nuttalliana*, all target samples amplified with a mean C_t_ = 24.18 for primer set AnuCOI1, and mean C_t_ = 23.49 for primer set AnuCOI2, and all non-target samples amplified at >11 C_t_ higher than targets for both assays. Because the primer set AnuCOI2 produced lower mean C_t_ values than primer set AnuCOI1, we proceeded with primer set AnuCOI2 for further testing. For *A. oregonensis*, all target samples amplified with a mean C_t_ = 24.98. All non-target samples either did not amplify or amplified at >12 C_t_ higher than targets. For all *Anodonta* primer sets, the melt curve produced a single sharp peak indicating no primer-dimer formation or off-target amplification. For *G. angulata*, all target samples amplified with all three primer sets, with mean Ct = 22.51 for primer set GanCOI1, mean C_t_ = 22.63 for GanCOI2, and mean C_t_ = 22.41 for GanCOI3. All non-targets either did not amplify or amplified at > 10 C_t_ higher than targets for all three primer sets, but primer set GanCOI2 had the greatest specificity, with all non-targets amplifying >12 C_t_ higher than targets. In addition, primer sets GanCOI1 and GanCOI3 displayed weak primer dimer formation in melt-curve analyses, while primer set GanCOI2 did not. Thus, primer set GanCOI2 was chosen for further work.

The *in vitro* evaluation of assay specificity including species-specific TaqMan™ MGB probes indicated that each assay was specific, detecting DNA from only targets species. For *A. nuttalliana*, all target samples amplified with both probes, with a mean C_t_ of 29.68 for probe AnuCOI2-Pa, and mean C_t_ 28.18 for probe AnuCOI2-Pb. No amplification was observed in any non-target samples for either probe. Because probe AnuCOI2-Pb displayed a lower mean C_t_ value than probe AnuCOI2-Pa, we proceeded with probe AnuCOI2-Pb for sensitivity testing and field validation. For *A. oregonensis* all target samples amplified with both probes with a mean C_t_ of 29.31 for probe AorCOI-Pa, and mean C_t_ 29.09 for probe AorCOI-Pb. No amplification was observed in any non-target samples for either probe. Because probe AorCOI-Pb displayed a lower mean C_t_ value than probe AorCOI-Pa, we proceeded with probe AorCOI-Pb for sensitivity testing and field validation. For *G. angulata*, all target samples amplified with probe GanCOI2-P with a mean C_t_ of 27.87, and no amplification was observed for any non-target samples.

Furthermore, the *A. nuttalliana* probe modified for use in the Central Valley of CA (AnuCOI2-Pc) amplified robustly with tissue DNA extracts from sites in the Pajaro, Sacramento, and San Joaquin Rivers. DNA extracts from ID, OR and UT also amplified robustly with both probes AnuCOI2-Pb and AnuCOI2-Pc combined at 125 nM each (Table 2). Thus, for work in the Central Valley of CA, we recommend using both probes in combination if it is not known which haplotype is found in a particular water body. Due to observed polymorphisms in central CA populations, if eDNA sampling is conducted in water bodies other than those for which sequence data was available for this study (Table S1), we recommend collecting and sequencing additional local tissue samples to ensure the optimal eDNA assay is used.

### Sensitivity testing

With the *A. nuttalliana/A. oregonensis* multiplex, for *A. nuttalliana*, all replicates amplified down to a template concentration of 5 copies per reaction, and 70% (7 of 10) of reactions containing 2 copies per reaction amplified in qPCR. For *A. oregonensis*, all replicates amplified down to 5 copies per reaction, and 40% (4 of 10) of reactions containing 2 copies per reaction amplified. With the *G. angulata/M. falcata* multiplex, for *G. angulata*, all replicates amplified down to 10 copies per reaction, 80% (8 of 10) reactions with 5 copies amplified, and 90% (9 of 10) reactions with 2 copies amplified. For *M. falcata*, all replicates amplified down to 10 copies per reaction, 60% (6 of 10) reactions with 5 copies amplified, and 30% (3 of 10) reactions with 2 copies amplified. With the *A. nuttalliana/M. falcata* multiplex, for *A. nuttalliana*, all replicates amplified down to 5 copies per reaction, and 70% (7 of 10) of reactions containing 2 copies per reaction amplified. For *M. falcata*, all replicates amplified down to 10 copies per reaction, 70% (7 of 10) reactions with 5 copies amplified, and 30% (3 of 10) reactions with 2 copies amplified.

### Field validation

In field validation with the *Anodonta* multiplex, we detected *A. nuttalliana* in 25/70 eDNA samples from 17/36 water bodies in 5 states, and *A. oregonensis* in 13/70 samples from 7/36 water bodies in 2 states (Table S4, Figure 2). Both *Anodonta* species were detected at 2 locations: Columbia Slough, OR, and Walla Walla River just above its confluence with the Columbia River, WA. Although traditional sampling was not conducted at all locations where eDNA samples were collected, live *Anodonta* were observed at 16 sites either at the time of, or within one year of eDNA sampling. At all 16 sites where live *Anodonta* were encountered, the corresponding eDNA sample was positive for one or both *Anodonta* species except for one site: the Chehalis River in WA (Table S4). At the Chehalis site, live *Anodonta* were found approximately 50 m downstream of where the eDNA sample was collected but *Anodonta* eDNA was not detected.

**Figure 2.**
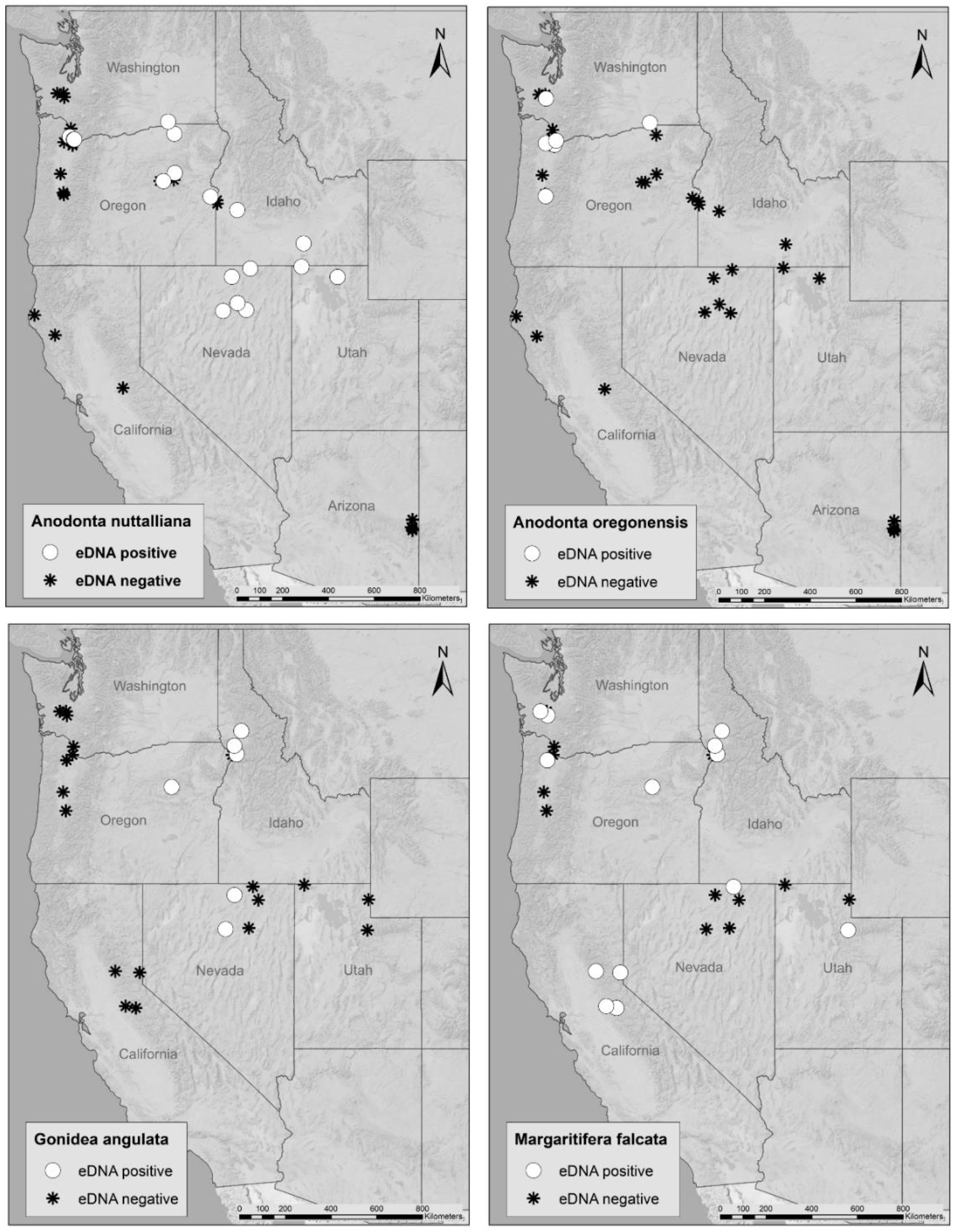
Locations of filtered water samples collected in the western United States for field validation of multiplex environmental DNA assays for *Anodonta nuttalliana, Anodonta oregonensis, Gonidea angulata* and *Margaritifera falcata.*

For the *G. angulata/M. falcata* multiplex, *G. angulata* were detected in 6/31 eDNA samples from 5/24 water bodies in 3 states, and *M. falcata* were detected in 18/31 samples from 13/24 water bodies in 6 states (Table S5, Figure 2). Live *G. angulata* individuals were encountered at 5 locations were eDNA samples were collected, and 100% of these eDNA samples were positive for *G. angulata.* Live *M. falcata* were encountered at 15 locations where eDNA samples were collected, and 100% of these eDNA samples were positive for *M. falcata* (Table S5).

For the *A. nuttalliana/M. falcata* multiplex, *A. nuttalliana* was detected in 13/29 eDNA samples, and *M. falcata* was detected in 18/29 samples (Table S6). In 2 samples, the Middle Fork John Day River, OR, and the Bruneau River, NV, both species were detected. Detection with the A. *nuttalliana/M. falcata* multiplex assay was 100% in agreement with detections of these species using the other two multiplex assays. No amplification was observed in any field blank, extraction blank, or qPCR negative control samples.

## Discussion

We designed and validated sensitive, species-specific qPCR assays for three freshwater mussel species (*A. nuttalliana*, *A. oregonensis*, and *G. angulata*) native to western North America, and we validated these in 3 multiplex assays along with a previously designed assay for *M. falcata* (Dysthe et al. 2018). These assays performed well for detecting mussels in the field at locations where they occur. Currently, *A. nuttalliana* and *G. angulata* are listed as vulnerable by the IUCN (Blevins et al. 2016a, 2016b), and some US states have recently listed them as ‘species of conservation concern’. Additionally, *G. angulata* has been assessed as endangered in Canada (COSEWIC 2010) and is listed as a species of special concern under Canada’s Species at Risk Act, and efforts are underway to petition listing of *G. angulata* for protection under the US Endangered Species Act (Blevins et al. *in prep*). *M. falcata* is considered near threatened and *A. oregonensis* is considered of least concern by the IUCN (Blevins et al. 2016c, 2016d), however dramatic declines have occurred in portions of the ranges of both species (Blevins et al. 2017), and they are also designated as ‘species of concern’ by some state agencies. Continued monitoring of these species will be essential to document persistence and to locate extant populations for protection.

When used in tandem, two of our multiplex assays can detect and discern all four western freshwater mussel species. In addition, we designed a third multiplex assay for use in areas where *A. nuttalliana* and *M. falcata* co-occur, but the other two species are not expected, likely a common scenario since these two species have the widest distributions of the four. It should be possible to use different multiplex permutations for specific sampling needs, although the three multiplex assays we designed and tested are likely sufficient for most sampling needs. It may be possible to multiplex all four assays using four different fluorescent dyes, but this would rarely be necessary or cost-effective because, to our knowledge, there are no sites where all 4 species co-occur, and only in a few locations (e.g., Middle Fork John Day River) do any three co-occur.

The assays described here will be particularly useful for assessing presence of freshwater mussel populations in locations with suitable habitat, or where populations have been previously described, with costs far lower than traditional stream or river-bed surveys (Stoeckle et al. 2016, Togaki et al. 2019). Further, our assays could be used in systematic monitoring programs to detect life history events such as glochidial releases or mussels die-offs, both of which would be expected to show dramatic increases in eDNA concentrations.

Aside from eDNA purposes, our assays may be useful for other applications. For example, these assays could be used for species identification of glochidia, the microscopic immature life stage of freshwater mussels, which are notoriously difficult to identify to species, typically requiring highly trained visual identification expertise or costly barcode sequencing. qPCR could potentially provide a more accurate and cost-effective solution for glochidia species identification. Because mussel glochidia attach to gills and/or fins of infected host fish, identifying glochidia could also be used to study species-specific host fish/mussel associations, which could have important implications for mussel transformation and dispersal (Strayer 2008, Maine et al. 2016).

Our *A. nuttalliana/A. oregonensis* qPCR assay could also provide low-cost species identification from mature mussel tissue samples. *A. nuttalliana* and *A. oregonensis* are morphologically similar and are often mis-identified for one another based on shell shape, and thus DNA barcode sequencing is often necessary to properly differentiate these species. Consequently, many historical records based on shell morphology of *A. nuttalliana* and *A. oregonensis* may be incorrect identifications. For example, some historical records indicate that *A. oregonensis* occurred in NV and UT, but neither recent genetic tissue analysis nor re-measurement of museum specimens supports *A. oregonensis* presence in NV or UT (Blevins et al. 2016b, Blevins et al. 2017). Likewise, we have not detected *A. oregonensis* in UT or NV with eDNA. Our qPCR assays could be a more cost-effective method of *Anodonta* species identification from tissue samples, and our multiplex assay for *Anodonta* will be useful for resampling historical *Anodonta* sites with eDNA to determine if original species identifications were correct.

One particular aspect of our eDNA assay design is more broadly informative for design of species-specific eDNA assays for other species. For *A. nuttalliana*, we were not able to design a single species-specific assay that was a perfect match to all range-wide haplotypes, likely because *A. nuttalliana* has a wide geographic distribution, and relatively high mitochondrial haplotype diversity due to prehistoric isolation in distinct hydrologic basins (Mock et al. 2010), while the other three western mussel species display lower levels of mitochondrial diversity (Mock et al. 2010, 2013). For *A. nuttalliana* we also had reference sequence data from a very large number of samples for assay design: 344 samples from 67 populations from 7 states. Thus, we likely uncovered sequence diversity that may have been missed had we designed the assay based on a more limited reference sequence library. Many published eDNA assays are based on far more limited reference sequence data that may not always account for rare haplotypes, which could affect assay sensitivity in certain populations.

Beyond the need to improve reference databases, future work should focus on establishing best practices for effectively sampling freshwater mussels with eDNA in the field. For example, further research is needed to determine transport distances of eDNA from mussel beds, and the degree to which they are modulated by physical and chemical water properties, to inform the spatial scale of sampling to ensure populations are not missed (eg. Deiner and Altermatt 2014, Wacker et al. 2019, Gasparini et al. 2020). Research to determine the optimal sampling season for maximum detection probability will also be important. Western freshwater mussels generally release millions of glochidia into the water column during the spring for dispersal by host fish (Culp et al. 2011, Allard et al. 2017), and these releases may provide an abundance of eDNA for detection at that time (Wacker et al. 2019). Alternatively, late summer through winter low flows may be efficacious for sampling because high water levels in spring can dilute eDNA, making it harder to detect.

As native mussel populations continue to decline throughout North America, efficient and reliable tools for monitoring them across large spatial and temporal scales are important for targeting conservation efforts. The multiplex eDNA assays described here will serve as valuable tools for future surveys of western freshwater mussels and will be more cost-effective than traditional sampling techniques. Additionally, they provide an effective means for distinguishing species that can be difficult to identify morphologically. This in turn will aid in the detection, monitoring, and conservation of native freshwater mussels across western North America.

## Acknowledgements

We would like to thank Krissy Wilson from the Utah Division of Wildlife Resources for seeing the value of using eDNA for western freshwater mussels, and for helping fund this study. We would also like to thank Kellie Carim and Yvette Paroz from the United States Forest Service, Alexa Maine from the Confederated Tribes of the Umatilla Indian Reservation, Emilie Blevins from the Xerces Society, Joel Sauder from Idaho Department of Fish and Game, and Christine O’Brien from Browns River Consultants for their advice and help with field work. We also thank Erika Rubenson, Zane Ruddy, Melissa Brown, and Jeff Sorenson for the original collection and repurposing of their eDNA samples. We acknowledge tissue samples obtained through the Confederated Tribes of the Umatilla Indian Reservation (Jayne Brim-Box, Donna Nez, David Wolf and Gene Shippentower), The Nature Conservancy (Jeanette Howard, Michelle Steg-Geltner), Washington Department of Fish and Wildlife (Molly Hallock), United States Geological Survey (Jason Dunham), Montana Natural Heritage Program (David Stagliano), Dan McGuire, Maria Ellis, and Lorrie Haley.

## Author Contributions

TWR conducted most laboratory and field work and wrote the initial manuscript draft. JCD and TWF assisted with assay design and validation and eDNA repurposing. JCD conducted some laboratory work, and CT aided with some field work. KEM, CT, JCD, TWF, and MKS provided edits and comments to the manuscript, and aided with study design.

