## supplementary Table S1 for "Detection of four imperiled western North American freshwater mussel species from environmental DNA with multiplex qPCR assays"

**Supplementary Information**

**Table S1**. Tissue samples used for designing quantitative PCR assays for detection of *Anodonta nuttalliana*, *Anodonta oregonensis* and *Gonidea angulata* from environmental DNA.

| *Anodonta nuttalliana* | | | | | | | |  |
| --- | --- | --- | --- | --- | --- | --- | --- | --- |
| Site | **# samples** | **Date Collected** | **State** | **County** | **Water body** | **Lat** | **Long** | **GenBank Accession #s** |
| ABR | 5 | 7/13/2004 | Arizona | Apache | Black River | 33.93635 | -109.40563 | MN912998 |
| ADY | 10 | 8/1/2009 | California | Tehama | Dye Creek | 40.10048 | -122.05449 | MN913005-MN913007 |
| AEA | 8 | 5/26/2001 | California | Mendocino | Eel River | 40.13911 | -123.81601 | MN913008, MN913008 |
| AEC | 10 | 7/17/2009 | California | Mendocino | S Fork Eel River | 39.74761 | -123.63330 | MN913010-MN913012 |
| AEH | 5 | 7/18/2009 | California | Humboldt | S Fork Eel River | 40.27236 | -123.85860 | MN913013, MN913014 |
| APB | 5 | 8/9/1995 | California | Shasta | Pit River | 40.98041 | -121.55817 | MN913043, MN913044 |
| APJ | 2 | 8/29/2009 | California | Santa Cruz | Pajaro River | 36.89405 | -121.64400 | MN913045 |
| APM | 5 | 8/15/2007 | California | Shasta | Pit River | 40.96497 | -121.79028 | MN913047 |
| APW | 10 | 12/15/2008 | California | Shasta | Pit River | 40.97846 | -121.54656 | MN913052, MN913053 |
| ARO | 9 | 7/25/2009 | California | Sonoma | Russian River | 38.47910 | -122.98984 | MN913054-MN913058 |
| ASA | 2 | 8/1/2009 | California | Glenn/Butte | Sacramento River | 39.62869 | -121.99345 | MN913062, MN913063 |
| ASY | 1 | 7/6/2008 | California | Siskiyou | Scott River | 41.63589 | -123.06951 | MN913089 |
| ATU | 10 | 10/21/2008 | California | Shasta | Tule River | 41.07987 | -121.44604 | MN913090 |
| AWD | 5 | 8/14/2007 | California | Plumas | Willow Creek Channel | 40.56262 | -120.63994 | MN913099 |
| AEW | 10 | 5/19/2007 | California | Mono | E Walker River | 38.44339 | -119.30656 | MN913015 |
| AOR | 10 | 5/18/2007 | California | Bishop | Owens River at Bishop Creek Canal | 37.40815 | -118.44062 | MN913037-MN913041 |
| AWE | 3 | 8/4/2009 | California | Mono | W. Fork Walker River | 38.61070 | -119.51775 | MN913100, MN913101 |
| APT | 10 | 8/19/2008 | California | Shasta | Pit River | 41.00028 | -121.75951 | MN913049-MN913051 |
| ASJ | 3 | 10/19/2008 | California | San Joaquin | San Joaquin River | 37.95127 | -121.33579 | MN913067, MN913068 |
| ASM | 10 | 10/7/2009 | California | San Joaquin | San Joaquin River | 37.95075 | -121.36133 | MN913069-MN913075 |
| ASV | 10 | 10/9/2009 | California | San Joaquin | San Joaquin River | 37.95104 | -121.32661 | MN913081-MN913087 |
| ASQ | 5 | 10/1/2009 | California | San Joaquin | San Joaquin River | 37.97317 | -121.37705 | MN913076-MN913078 |
| ASF | 2 | 7/16/2007 | Idaho | Elmore | Snake River | 42.93847 | -115.31987 | MN913064, MN913065 |
| AWP | 9 | 9/2/2002 | Idaho | Power | Lake Walcott | 42.62585 | -113.10608 | MN913107, MN913108 |
| ABY | 1 | 7/24/2008 | Idaho | Owyhee | Bruneau River | Missing | missing | MN913110 |
| ARR | 1 | 8/23/2006 | Nevada | Lander | Upper Reese River | 39.18682 | -117.34831 | MN913059 |
| ARU | 1 | 8/9/2007 | Nevada | Lander | Upper Reese River | 39.19012 | -117.34809 | MN913061 |
| ABB | 2 | 8/16/2004 | Oregon | Multnomah | Bybee Lake | 45.63232 | -122.68529 | MN912995 |
| ACC | 6 | 6/11/2005 | Oregon | Multnomah | Columbia | 45.56301 | -122.53488 | MN913000, MN913001 |
| ACT | 9 | 6/8/2005 | Oregon | Deschutes | Crooked River | 44.36641 | -121.14096 | MN913002 |
| ADC | 11 | 6/9/2005 | Oregon | Wasco | Deschutes River | 45.62934 | -120.91085 | MN913004 |
| AWW | 8 | 6/7/2005 | Oregon | Linn | Willamette River | 44.45399 | -123.20998 | MN913109 |
| ACU | 6 | 7/7/2003 | Oregon | Umatilla | Umatilla | 45.83506 | -119.33269 | MN913003 |
| AFH | 5 | 8/2/2007 | Oregon | Grant | Middle Fork John Day | 44.79940 | -118.97629 | MN913016, MN913017 |
| AJC | 2 | 6/11/2003 | Oregon | Grant | Middle Fork John Day | 44.69361 | -118.79746 | MN913018 |
| AJF | 7 | 6/5/2005 | Oregon | Grant | Middle Fork John Day | 44.75985 | -118.86547 | MN913019-MN913021 |
| AJH | 5 | 9/28/2005 | Oregon | Grant | Middle Fork John Day | 44.75985 | -118.86547 | MN913022-MN913024 |
| AJR | 5 | 8/4/2007 | Oregon | Grant | Middle Fork John Day | 44.88807 | -119.19683 | MN913025-MN913027 |
| AJS | 8 | 6/11/2003 | Oregon | Grant | Middle Fork John Day | 44.75985 | -118.86547 | MN913028, MN913029 |
| AJW | 5 | 9/28/2005 | Oregon | Grant | Middle Fork John Day | 44.79901 | -118.93652 | MN913030-MN913032 |
| AMI | 5 | 9/10/2004 | Oregon | Malheur | McDermitt Creek | 42.02007 | -117.93437 | MN913035 |
| AOT | 3 | 9/6/2004 | Idaho | Owyhee | Owyhee | 42.57574 | -117.00174 | MN913042 |
| AUF | 1 | 7/7/2003 | Oregon | Umatilla | Umatilla | 45.86733 | -119.31858 | MN913091 |
| AUH | 5 | 6/2/2005 | Oregon | Umatilla | Umatilla River | 45.79195 | -119.32155 | MN913092-MN913093 |
| AUW | 4 | 7/8/2003 | Oregon | Umatilla | Umatilla | 45.80371 | -119.36037 | MN913095-MN913097 |
| AWF | 5 | 7/20/2007 | Oregon | Walla Walla | Walla Walla River | 46.06219 | -118.90361 | MN913102-MN913104 |
| AWH | 5 | 6/7/2003 | Oregon | Umatilla | Umatilla | 45.74507 | -118.59963 | MN913105 |
| AWM | 5 | 9/27/2007 | Oregon | Umatilla | Wildhorse Creek | 45.72161 | -118.67208 | MN913106 |
| APL | 5 | 8/21/2001 | Utah | Millard | Pruess Lake | 38.89871 | -114.00415 | MN913046 |
| ASG | 1 | 9/29/2003 | Utah | Box Elder | Snake River | 41.99002 | -113.99459 | MN913066 |
| ABP | 5 | 8/22/2001 | Utah | Juab | Burriston Pond | 39.79762 | -111.86892 | MN912996, MN912997 |
| ABU | 5 | 8/17/2001 | Utah | Rich | Bear River | 41.53697 | -111.13429 | MN912999 |
| AOC | 5 | 8/22/2001 | Utah | Piute | Otter Creek Reservoir | 38.16946 | -112.02215 | MN913036 |
| APR | 5 | 8/22/2001 | Utah | Piute | Piute Reservoir | 38.25306 | -112.21369 | MN913048 |
| ARS | 5 | 8/21/2001 | Utah | Tooele | Redden Springs | 40.01139 | -113.74158 | MN913060 |
| ASU | 1 | 8/14/2001 | Utah | Box Elder | Salt Creek | 41.71497 | -112.23016 | MN913080 |
| AWC | 5 | 6/5/2007 | Washington | Yakima | Wenas Creek | 46.69584 | -120.49408 | MN913098 |
| AKR | 2 | 7/25/2007 | Washington | Ferry | Kettle River | 48.88386 | -118.60111 | MN913033 |
| ALC | 5 | 7/23/2007 | Washington | Lincoln | Lords Creek | 47.38655 | -118.14368 | MN913034 |
| AST | 5 | 7/24/2007 | Washington | Asotin | Snake River | 46.56309 | -118.10046 | MN913079 |
| ASW | 3 | 5/7/2007 | Washington | Loomis | Sinlahekin Creek | 48.75581 | -119.65780 | MN913088 |
| AHL | 3 | 9/5/2017 | Nevada | Elko | Humboldt | 40.69139 | -115.83430 | MN913111 |
| AHU | 3 | 9/5/2017 | Nevada | Elko | Humboldt | 40.63479 | -115.73650 | MN913112, MN913113 |
| AMC | 3 | 9/5/2017 | Nevada | Eureka | Humboldt | 40.89337 | -116.17617 | MN913115, MN913116 |
| AUB | 3 | 9/6/2017 | Nevada | Elko | Bruneau | 41.52487 | -115.46136 | MN913119-MN913121 |
| ALB | 3 | 9/6/2017 | Nevada | Elko | Bruneau | 41.67072 | -116.39766 | MN913114 |
| ASO | 3 | 9/7/2017 | Nevada | Elko | Owyhee | 41.67072 | -116.39766 | MN913117, MN913118 |
| *Anodonta oregonensis* | | | | | | | | |
| ANC | 3 | 6/7/2007 | Alaska | Prince of Wales Island | Naukati Creek | 55.90867 | -133.09917 | MN945935 |
| AHY | 2 | 6/14/2007 | British Columbia | na | Hathaway Lake | 51.66666 | -120.83335 | MN945922 |
| AKCR | 8 | 5/17/2006 | British Columbia | na | Lake Chilliwack | 49.08687 | -121.45252 | MN945923, MN945924 |
| AKSB | 1 | 7/7/2005 | British Columbia | na | Slocan River | 49.67950 | -117.65700 | MN945929 |
| AKD | 5 | 5/1/2008 | California | Siskiyou | Klamath River | 41.93090 | -122.44229 | MN945925-MN945927 |
| ALR | 2 | 8/3/2009 | California | Siskiyou | Lost River | 41.99820 | -121.52209 | MN945931, MN945932 |
| API | 4 | 8/3/2009 | California | Modoc | Pit River | 41.40058 | -120.93422 | MN945936 |
| AAF | 5 | 10/1/2003 | Oregon | Lake | Summer Lake | 42.99226 | -120.74321 | MN945908 |
| ABB | 9 | 8/16/2004 | Oregon | Multnomah | Bybee Lake | 45.63232 | -122.68529 | MN945909-MN945912 |
| ACC | 4 | 6/11/2005 | Oregon | Multnomah | Columbia | 45.56301 | -122.53488 | MN945913-MN945915 |
| ACS | 5 | 10/1/2003 | Oregon | Lake | Coyote Creek | 42.85599 | -121.16044 | MN945916, MN945917 |
| ASL | 5 | 10/1/2003 | Oregon | Lake County | Sycan River | 42.77994 | -121.11420 | MN945938-MN945940 |
| AWW | 3 | 6/7/2005 | Oregon | Linn | Willamette River | 44.45399 | -123.20998 | MN945946 |
| ACU | 4 | 7/7/2003 | Oregon | Umatilla | Columbia | 45.83506 | -119.33269 | MN945918 |
| AUH | 5 | 6/2/2005 | Oregon | Umatilla | Umatilla River | 45.79195 | -119.32155 | MN945942, MN945943 |
| AWF | 5 | 7/20/2007 | Oregon | Walla Walla | Walla Walla River | 46.06219 | -118.90361 | MN945944, MN945945 |
| AKH | 1 | 2/8/1993 | Oregon | missing | Klamath River | missing | missing | MN945928 |
| ACV | 5 | 8/8/2007 | Washington | Grays Harbor | Chehalis River | 46.98187 | -123.47097 | MN945919 |
| ALD | 2 | 9/1/2008 | Washington | Clallam | Lake Sutherland | 48.08066 | -123.72176 | MN945930 |
| AML | 5 | 8/29/2007 | Washington | Mason | Mason Lake | 47.351337 | -122.92200 | MN945933, MN945934 |
| AHS | 5 | 9/16/2007 | Washington | Benton | Handford Slough | 46.58917 | -119.38223 | MN945920, MN945921 |
| ASK | 2 | 9/30/2006 | Washington | Okanogan | Similkameen River | 48.95341 | -119.64872 | MN945937 |
| ASW | 5 | 5/7/2007 | Washington | Loomis | Sinlahekin Creek | 48.75581 | -119.65780 | MN945941 |
| *Gonidea angulata* | | | | | | | | |
| GJB | 3 | 6/11/2003 | Oregon | Grant | Middle Fork John Day | 44.75984982 | -118.8654736 | MN957813, MN957814 |
| GJF | 5 | 6/3/2005 | Oregon | Grant | Middle Fork John Day | 44.73855141 | -118.8476993 | MN957815-MN957817 |
| GJW | 4 | 9/28/2005 | Oregon | Grant | Middle Fork John Day | 44.79900587 | -118.9365159 | MN957824, MN957825 |
| GJH | 5 | 9/28/2005 | Oregon | Grant | Middle Fork John Day | 44.75984982 | -118.8654736 | MN957818-MN957820 |
| GUF | 3 | 7/7/2003 | Oregon | Umatilla | Umatilla River | 45.86732814 | -119.3185774 | MN957863, MN957864 |
| GUD | 1 | 7/8/2003 | Oregon | Umatilla | Umatilla River | 45.79374016 | -119.3268452 | MN957862 |
| GOB | 5 | 9/11/2004 | Oregon | Malheur | Owyhee River | 43.23043705 | -117.4994141 | MN957837-MN957840 |
| GFH | 5 | 8/2/2007 | Oregon | Grant | Middle Fork John Day | 44.79939546 | -118.9762937 | MN957808, MN957809 |
| GJR | 5 | 8/4/2007 | Oregon | Grant | Middle Fork John Day | 44.88806838 | -119.1968323 | MN957821-MN957823 |
| GCE | 1 | 8/2/2004 | Washington | Grays Harbor | Chehalis River | 46.98153761 | -123.4124825 | MN957802 |
| GCV | 5 | 8/8/2007 | Washington | Grays Harbor | Chehalis River | 46.98187889 | -123.4709721 | MN957806, MN957807 |
| GTC | 5 | 9/9/2005 | Washington | Yakima | Toppenish Creek | 46.30920706 | -120.3312899 | MN957859-MN957861 |
| GSK | 5 | 9/31/2006 | Washington | Okanogan | Similkameen River | 48.95341054 | -119.6487164 | MN957846, MN957847 |
| GSO | 5 | 7/25/2007 | Washington | Asotin | Snake River | 46.16227086 | -116.9232958 | MN957848-MN957851 |
| GPM | 5 | 8/15/2007 | California | Shasta | Pit River | 40.96497028 | -121.7902827 | MN957843, MN957844 |
| GKC | 5 | 6/13/2007 | California | Siskiyou | Klamath River | 41.798297 | -123.316786 | MN957826, MN957828 |
| GSY | 5 | 7/6/2008 | California | Siskiyou | Scott River | 41.63588898 | -123.0695058 | MN957856-MN957858 |
| GKF | 5 | 7/7/2008 | California | Siskiyou | Klamath River | 41.72297977 | -123.4370735 | MN957829, MN957830 |
| GKK | 5 | 7/7/2008 | California | Siskiyou | Klamath River | 41.71052298 | -123.448757 | MN957831-MN957833 |
| GKT | 5 | 7/8/2008 | California | Siskiyou | Klamath River | 41.78809185 | -123.0513288 | MN957836 |
| GKL | 5 | 7/8/2008 | California | Siskiyou | Klamath River | 41.85577581 | -122.57177 | MN957834, MN957835 |
| GSB | 10 | 7/9/2008 | California | Siskiyou | Shasta River | 41.80690236 | -122.5935359 | MN957845 |
| GHR | 5 | 9/5/2017 | Nevada | Lander | Humboldt | 40.661208 | -116.746701 | MN957810-MN957812 |
| GOS | 5 | 9/7/2017 | Nevada | Elko | S. Fork Owyhee | 41.67072078 | -116.3976579 | MN957841, MN957842 |
| GSS | 7 | 8/15/2018 | Idaho | Idaho | Salmon River | 45.90180 | -116.40460 | MN957852-MN957855 |
| GCP | 5 | 8/15/2018 | Idaho | Clearwater | Clearwater River | 46.29880 | -116.12800 | MN957803-MN957805 |
| OKV | 6 | 5/21/2016 | British Columbia |  | Vaseux lake | 49.301025 | -119.531299 | MN957868-MN957870 |
| RLC | 5 | 8/26/2016 | British Columbia |  | Okanagan River | 49.043821 | -119.468089 | MN957875-MN957878 |
| RFV | 5 | 8/26/2016 | British Columbia |  | Okanagan River | 49.192434 | -119.550677 | MN957873, MN957874 |
| LKB | 5 | 9/12/2016 | British Columbia |  | S. Okanagan Lake | 49.60754 | -119.650085 | MN957866, MN957867 |
| KKB | 5 | 8/31/2016 | British Columbia |  | N. Okanagan Lake | 50.250238 | -119.350013 | MN957865 |

**Table S2.** Sequences retrieved from GenBank used to design quantitative PCR assays for the detection *of Anodonta nuttalliana, Anodonta oregonensis* and *Gonidea Angulata* from environmental DNA.

| *Anodonta nuttalliana* | |
| --- | --- |
| Accession number | **Location** |
| DQ272367.1 | Kalama WA |
| DQ272365.1 | Kalama WA |
| DQ272369.1 | Kalama WA |
| DQ272368.1 | Kalama WA |
| DQ272366.1 | Kalama WA |
| KX822634.1 | Unknown (USA) |
| EU327355.1 | Haplotype attributed to samples from various locations |
| KF672864.1 | Black River, AZ |
| KF672865.1 | Arroyo de la Concha, Mexico |
| KF672866.1 | Rio Tres Rios, Mexico |
| KF672867.1 | Rio La Palotada, Mexico |
| KF672868.1 | Rio La Palotada, Mexico |
| KF672869.1 | Black River, AZ |
| KF672870.1 | Rio La Palotada, Mexico |
| KF672871.1 | Rio Santa Maria, Mexico |
| KF672872.1 | Rio Tres Rios, Mexico |
| KF672873.1 | Rio Tres Rios, Mexico |
| KF672875.1 | Arroyo de la Concha, Mexico |
| KF672876.1 | Rio La Palotada, Mexico |
| KF672877.1 | Black River, AZ |
| KF672878.1 | Black River, AZ |
| KF672879.1 | Rio Tres Rios, Mexico |
| KF672882.1 | Owens River, CA |
| KF672891.1 | Owens River, CA |
| KF672892.1 | Owens River, CA |
| KF672893.1 | Owens River, CA |
| EU327356.1 | Haplotype attributed to samples from various locations |
| Anodonta oregonensis | |
| DQ272359.1 | Ozette WA |
| DQ272360.1 | Sand point WA |
| DQ272361.1 | Pleasure Point WA |
| DQ272362.1 | Lake Sammamish WA |
| DQ272363.1 | Seward Park WA |
| DQ272364.1 | Seward Park WA |
| EU327354.1 | Haplotype attributed to samples from various locations |
| EU327353.1 | Haplotype attributed to samples from various locations |
| EU327352.1 | Haplotype attributed to samples from various locations |
| EU327350.1 | Haplotype attributed to samples from various locations |
| EU327351.1 | Haplotype attributed to samples from various locations |
| Gonidea Angulata | |
| DQ272373.1 | Greeley Bar OR |
| DQ272372.1 | Greeley Bar OR |
| DQ272371.1 | Greeley Bar OR |
| AF231755.1 | Unknown |
| KP795030.1 | Unknown |
| DQ191412.1 | Unknown |

**Table S3.** GenBank derived sequences from non-target species used to design quantitative PCR assays for the detection of *Anodonta nuttalliana, Anodonta oregonensis* and *Gonidea angulata* from environmental DNA.

| Species | GenBank Accession # |
| --- | --- |
| *Anodonta beringiana* | DQ272370.1 |
| *Anodonta beringiana* | EU327357.1 |
| *Corbicula fluminea* | KT893358.1 |
| *Corbicula fluminea* | KT893360.1 |
| *Corbicula fluminea* | DQ264393.1 |
| *Corbicula fluminea* | KT893330.1 |
| *Corbicula fluminea* | KT893331.1 |
| *Corbicula fluminea* | KT893333.1 |
| *Corbicula fluminea* | KT893354.1 |
| *Dreissena polymorpha* | JQ435817.1 |
| *Dreissena polymorpha* | DQ840124.1 |
| *Dreissena polymorpha* | U47653.1 |
| *Dreissena polymorpha* | AF120663.1 |
| *Dreissena polymorpha* | EU484456.1 |
| *Dreissena bugensis* | U47651.1 |
| *Dreissena bugensis* | EU484436.1 |
| *Dreissena bugensis* | DQ840132.1 |
| *Ferrissia rivularis* | KF737915.1 |
| *Ferrissia rivularis* | KF737914.1 |
| *Ferrissia rivularis* | KF737913.1 |
| *Lampsilis siliquoidea* | KC408781.1 |
| *Lampsilis siliquoidea* | KC408744.1 |
| *Lampsilis siliquoidea* | KC408772.1 |

**Table S4.** Locations of eDNA samples collected for detection of *A. nuttalliana* and *A. oregonensis*. All samples were run in qPCR with a multiplex eDNA assay designed for detecting and discerning both *Anodonta* species.

| River/Stream | State | Latitude | Longitude | Collection date | A. nuttalliana eDNA detected (Y/N) | A. oregonensis eDNA detected (Y/N) | Live Anodonta encountered (Y/N) |
| --- | --- | --- | --- | --- | --- | --- | --- |
| Boneyard Creek | AZ | 33.86535 | -109.30266 | 6/14/2017 | N | N | N |
| Boneyard Creek | AZ | 33.86665 | -109.29875 | 6/14/2017 | N | N | N |
| Boneyard Creek | AZ | 33.87069 | -109.29854 | 6/14/2017 | N | N | N |
| Boneyard Creek | AZ | 33.87216 | -109.29717 | 6/13/2017 | N | N | N |
| Boneyard Creek | AZ | 33.87846 | -109.28244 | 6/13/2017 | N | N | N |
| EF Black River | AZ | 33.81136 | -109.30668 | 6/20/2017 | N | N | N |
| EF Black River | AZ | 33.85323 | -109.31569 | 6/12/2017 | N | N | N |
| EF Black River | AZ | 33.85355 | -109.31543 | 6/17/2017 | N | N | N |
| EF Black River | AZ | 33.85405 | -109.31624 | 6/12/2017 | N | N | N |
| Becker Lake | AZ | 34.15181 | -109.30879 | 6/15/2017 | N | N | N |
| Little Colorado | AZ | 34.16243 | -109.30223 | 6/15/2017 | N | N | N |
| NF EF Black River | AZ | 33.91036 | -109.34713 | 6/14/2017 | N | N | N |
| NF EF Black River | AZ | 33.91174 | -109.34424 | 6/14/2017 | N | N | N |
| NF EF Black River | AZ | 33.91175 | -109.34508 | 6/14/2017 | N | N | N |
| NF Eel River | CA | 39.93682 | -123.34582 | 6/5/2017 | N | N | N |
| NF Eel River | CA | 39.96727 | -123.33838 | 6/5/2017 | N | N | N |
| NF Eel River | CA | 40.13487 | -123.35567 | 6/7/2017 | N | N | N |
| NF Eel River | CA | 40.13769 | -123.35587 | 6/7/2017 | N | N | N |
| Van Duzen River | CA | 40.53710 | -124.14730 | 6/12/2017 | N | N | N |
| Mokelumne River | CA | 38.32809 | -120.67710 | 7/10/2018 | N | N | N |
| Boise River | ID | 43.59945 | -116.18718 | 8/21/2017 | Y | N | N |
| Boise River | ID | 43.78153 | -116.97256 | 8/30/2017 | N | N | N |
| Malheur River | ID | 43.97781 | -117.23846 | 8/30/2017 | Y | N | N |
| Snake River | ID | 42.64179 | -113.57777 | 8/22/2017 | Y | N | N |
| Snake River | ID | 42.65304 | -113.56588 | 8/22/2017 | N | N | N |
| Snake River | ID | 42.65379 | -113.56304 | 8/22/2017 | N | N | N |
| Snake River | ID | 42.65379 | -113.56304 | 8/22/2017 | N | N | N |
| Snake River | ID | 43.87612 | -116.98460 | 8/30/2017 | N | N | N |
| SF Owyhee River | NV | 41.67072 | -116.39766 | 9/7/2017 | Y | N | Y |
| Bruneau River | NV | 41.91220 | -115.67523 | 9/6/2017 | Y | N | Y |
| Bruneau River | NV | 41.52487 | -115.46136 | 9/6/2017 | Y | N | Y |
| Humboldt River | NV | 40.66121 | -116.74670 | 9/5/2017 | Y | N | N |
| Maggie Creek | NV | 40.89337 | -116.17617 | 9/5/2017 | Y | N | Y |
| SF Humboldt River | NV | 40.63479 | -115.73650 | 9/5/2017 | Y | N | Y |
| SF Humboldt River | NV | 40.69139 | -115.83430 | 9/5/2017 | Y | N | Y |
| Crystal Springs Creek | OR | 45.46305 | -122.64270 | 1/26/2018 | N | Y | N |
| Crystal Springs Creek | OR | 45.47255 | -122.64220 | 1/26/2018 | N | Y | N |
| Crystal Springs Creek | OR | 45.47255 | -122.64218 | 10/12/2017 | N | Y | N |
| Crystal Springs Creek | OR | 45.47502 | -122.64170 | 1/26/2018 | N | Y | N |
| Crystal Springs Creek | OR | 45.47502 | -122.64173 | 10/13/2017 | N | Y | N |
| Crystal Springs Creek | OR | 45.48204 | -122.63709 | 10/13/2017 | N | Y | N |
| Crystal Springs Creek | OR | 45.48204 | -122.63710 | 1/26/2018 | N | Y | N |
| John Day River | OR | 44.41510 | -119.08578 | 7/22/2016 | Y | N | N |
| John Day River | OR | 44.41757 | -118.94579 | 7/22/2016 | Y | N | N |
| John Day River | OR | 44.41832 | -119.22548 | 7/22/2016 | N | N | N |
| John Day River | OR | 44.42060 | -118.97447 | 7/22/2016 | Y | N | N |
| John Day River | OR | 44.43576 | -119.29130 | 7/22/2016 | N | N | N |
| John Day River | OR | 44.44844 | -118.74613 | 7/22/2016 | Y | N | N |
| John Day River | OR | 44.45391 | -118.67779 | 7/22/2016 | N | N | N |
| MF John Day River | OR | 44.64156 | -118.63872 | 7/21/2016 | Y | N | N |
| MF John Day River | OR | 44.66855 | -118.71147 | 7/21/2016 | Y | N | N |
| MF John Day River | OR | 44.71736 | -118.82212 | 7/21/2016 | Y | N | N |
| Columbia Slough | OR | 45.57234 | -122.58533 | 9/26/2018 | Y | Y | Y |
| Calapooia River | OR | 44.61814 | -123.13049 | 9/25/2018 | N | N | N |
| Makenzie River | OR | 44.09087 | -123.02182 | 9/25/2018 | N | N | N |
| MF Williamette | OR | 44.02642 | -122.99652 | 9/25/2018 | N | Y | Y |
| Oswego Creek | OR | 45.41061 | -122.66199 | 9/24/2018 | N | N | N |
| Tualatin River | OR | 45.50015 | -122.99100 | 9/24/2018 | N | Y | N |
| Willamette River | OR | 45.65100 | -122.76298 | 9/24/2018 | Y | N | Y |
| Whitiker slough | OR | 45.57454 | -122.60921 | 9/24/2018 | Y | N | Y |
| MF John Day River | OR | 44.75984 | -118.86547 | 6/24/2016 | Y | N | Y |
| Wildhorse Creek | OR | 45.72721 | -118.65365 | 6/23/2016 | Y | N | Y |
| Raft River | UT | 41.96720 | -113.66016 | 6/28/2017 | Y | N | Y |
| Salt Creek | UT | 41.66362 | -112.24319 | 11/22/2016 | Y | N | Y |
| Lewis River | WA | 45.87209 | -122.72222 | 9/26/2018 | N | N | N |
| Scatter Creek | WA | 46.82856 | -123.00887 | 9/26/2018 | N | N | N |
| Chehalis River | WA | 46.83060 | -123.25783 | 9/26/2018 | N | N | Y |
| Fort Boerst lake | WA | 46.72374 | -122.97875 | 9/26/2018 | N | Y | N |
| Skookumchuck River | WA | 46.72124 | -122.97728 | 9/26/2018 | N | Y | N |
| Walla-Walla River | WA | 46.06301 | -118.90302 | 6/23/2016 | Y | Y | Y |

**Table S5.** Locations of eDNA samples collected for detection of *M. falcata* and *G. angulata*. All samples were run in qPCR with a multiplex eDNA assay designed for detecting and discerning both species.

| River/Stream | State | Latitude | Longitude | Collection date | M. falcata eDNA detected (Y/N) | G. angulata eDNA detected (Y/N) | Live M. falcata encountered (Y/N) | Live G. angulata encountered (Y/N) |
| --- | --- | --- | --- | --- | --- | --- | --- | --- |
| Oregon Creek | CA | 39.39537 | -121.08324 | 7/2/2018 | Y | N | Y | N |
| Oregon Creek | CA | 39.39589 | -121.08298 | 7/2/2018 | Y | N | Y | N |
| Oregon Creek | CA | 39.39540 | -121.08327 | 7/2/2018 | Y | N | Y | N |
| Oregon Creek | CA | 39.39512 | -121.08361 | 7/2/2018 | Y | N | Y | N |
| Yuba river | CA | 39.39461 | -121.08310 | 7/2/2018 | Y | N | Y | N |
| Truckee River | CA | 39.35260 | -120.12156 | 7/2/2018 | Y | N | Y | N |
| Stanislaus River | CA | 38.26971 | -120.27089 | 7/9/2018 | Y | N | Y | N |
| Mokelumne River | CA | 38.32809 | -120.67710 | 7/10/2018 | Y | N | Y | N |
| Snake River | ID | 45.64760 | -116.48670 | 8/15/2018 | N | N | N | N |
| Salmon River | ID | 45.66460 | -116.31250 | 8/15/2018 | Y | Y | Y | Y |
| Salmon River | ID | 45.90180 | -116.40460 | 8/15/2018 | Y | Y | N | N |
| Clearwater River | ID | 46.29880 | -116.12800 | 8/23/2018 | Y | Y | Y | Y |
| SF Humboldt River | NV | 40.69139 | -115.83430 | 9/5/2017 | N | N | N | N |
| SF Humboldt River | NV | 40.63479 | -115.73650 | 9/5/2017 | N | N | N | N |
| Humboldt river | NV | 40.66121 | -116.74670 | 9/5/2017 | N | Y | N | Y |
| Bruneau River | NV | 41.52487 | -115.46136 | 9/6/2017 | N | N | N | N |
| Bruneau River | NV | 41.91220 | -115.67523 | 9/6/2017 | Y | N | Y | N |
| SF Owyhee River | NV | 41.67072 | -116.39766 | 9/7/2017 | N | Y | N | Y |
| Middle Fork John Day | OR | 44.75984 | -118.86547 | 6/24/2016 | Y | Y | Y | Y |
| Willamette River | OR | 45.65100 | -122.76298 | 9/24/2018 | N | N | N | N |
| Tualatin River | OR | 45.50015 | -122.99100 | 9/24/2018 | Y | N | N | N |
| Makenzie River | OR | 44.09087 | -123.02182 | 9/25/2018 | N | N | N | N |
| Calapooia River | OR | 44.61814 | -123.13049 | 9/25/2018 | N | N | N | N |
| Bear River | UT | 41.53216 | -111.13196 | 11/15/2016 | N | N | N | N |
| Raft River | UT | 41.96720 | -113.66016 | 6/28/2017 | N | N | N | N |
| Beaver Creek | UT | 40.62783 | -111.17947 | 10/11/2017 | Y | N | Y | N |
| Beaver Creek | UT | 40.63082 | -111.18675 | 10/11/2017 | Y | N | Y | N |
| Skookumchuck River | WA | 46.72124 | -122.97728 | 9/26/2018 | Y | N | N | N |
| Chehalis River | WA | 46.83060 | -123.25783 | 9/26/2018 | Y | N | Y | N |
| Scatter Creek | WA | 46.82856 | -123.00887 | 9/26/2018 | N | N | N | N |
| Lewis River | WA | 45.87209 | -122.72222 | 9/26/2018 | N | N | N | N |

**Table S6.** Locations of eDNA samples collected for detection of *M. falcata* and *A. nuttalliana*. All samples were run in qPCR with a multiplex eDNA assay designed for detecting and discerning both species.

| River/Stream | State | Latitude | Longitude | Collection date | A. nuttalliana eDNA detected (Y/N) | M. falcata eDNA detected (Y/N) | Live A. nuttalliana encountered (Y/N) | Live M. falcata encountered (Y/N) |
| --- | --- | --- | --- | --- | --- | --- | --- | --- |
| Mokelumne River | CA | 38.32809 | -120.67710 | 7/10/2018 | N | Y | N | Y |
| Stanislaus River | CA | 38.26971 | -120.27089 | 7/9/2018 | N | Y | N | Y |
| Truckee River | CA | 39.35259 | -120.12150 | 7/2/2018 | N | Y | N | Y |
| Yuba river | CA | 39.39461 | -121.08310 | 7/2/2018 | N | Y | N | Y |
| Oregon Creek | CA | 39.39512 | -121.08361 | 7/2/2018 | N | Y | N | Y |
| Oregon Creek | CA | 39.39540 | -121.08327 | 7/2/2018 | N | Y | N | Y |
| Oregon Creek | CA | 39.39589 | -121.08298 | 7/2/2018 | N | Y | N | Y |
| Oregon Creek | CA | 39.39537 | -121.08324 | 7/2/2018 | N | Y | N | Y |
| Clearwater River | ID | 46.29880 | -116.12800 | 8/23/2018 | N | Y | N | Y |
| Salmon River | ID | 45.90180 | -116.40460 | 8/15/2018 | N | Y | N | Y |
| Salmon River | ID | 45.66460 | -116.31250 | 8/15/2018 | N | Y | N | Y |
| SF Owyhee River | NV | 41.67072 | -116.39766 | 9/7/2017 | Y | N | Y | N |
| Bruneau River | NV | 41.91220 | -115.67523 | 9/6/2017 | Y | Y | Y | Y |
| Bruneau River | NV | 41.52487 | -115.46136 | 9/6/2017 | Y | N | Y | N |
| Maggie Creek | NV | 40.89337 | -116.17617 | 9/5/2017 | Y | N | Y | N |
| SF Humboldt River | NV | 40.63479 | -115.73650 | 9/5/2017 | Y | N | Y | N |
| SF Humboldt River | NV | 40.69139 | -115.83430 | 9/5/2017 | Y | N | Y | N |
| Columbia Slough | OR | 45.57234 | -122.58533 | 9/26/2018 | Y | N | Y | N |
| Tualatin River | OR | 45.50015 | -122.99100 | 9/24/2018 | N | Y | N | N |
| willamette River | OR | 45.65100 | -122.76298 | 9/24/2018 | Y | N | Y | N |
| Whitiker slough | OR | 45.57454 | -122.60921 | 9/24/2018 | Y | N | Y | N |
| MF John Day | OR | 44.75984 | -118.86547 | 6/24/2016 | Y | Y | y | y |
| Wildhorse Creek | OR | 45.72745 | -118.65280 | 6/23/2016 | Y | N | Y | N |
| Beaver Creek | UT | 40.63082 | -111.18675 | 10/11/2017 | N | Y | N | Y |
| Beaver Creek | UT | 40.62783 | -111.17947 | 10/11/2017 | N | Y | N | Y |
| Raft River | UT | 41.96720 | -113.66016 | 6/28/2017 | Y | N | Y | N |
| Salt Creek | UT | 41.66362 | -112.24319 | 11/22/2016 | Y | N | Y | N |
| Chehalis River | WA | 46.83060 | -123.25783 | 9/26/2018 | N | Y | N | Y |
| Skookumchuck River | WA | 46.72124 | -122.97728 | 9/26/2018 | N | Y | N | Y |

**Table S7.** Species identified by NCBI primer blast that could potentially amplify with the AorCOI primer set. Primer specificity stringency settings were: Primer must have at least 2 total mismatches to unintended targets, including at least 1 mismatch within the last 4 bps at the 3' end. Ignore targets that have 5 or more mismatches.

| Species name | Range | Organism type |
| --- | --- | --- |
| *Lasmigona compressa* | NE US and SE Canada | Freshwater mussel |
| *Nodularia douglasiae* | Asia | Freshwater mussel |
| *Lasmigona subviridis* | US East Coast | Freshwater mussel |
| *Hemistena lata* | Alabama | Freshwater mussel |
| *Obovaria olivaria* | Eastern and Midwest US, eastern Canada | Freshwater mussel |
| *Epioblasma brevidens* | SE US | Freshwater mussel |
| *Leptodea fragilis* | Eastern and Midwest US, eastern Canada | Freshwater mussel |
| *Lasmigona decorata* | Carolinas | Freshwater mussel |
| *Alasmidonta heterodon* | Eastern US | Freshwater mussel |
| *Fusconaia mitchelli* | New Mexico and Texas | Freshwater mussel |
| *Potamilus amphichaenus* | Texas and Louisiana | Freshwater mussel |
| *Unio tumidiformis* | Europe | Freshwater mussel |
| *Tegula funebralis* | West Coast Marine | Marine mussel |
| *Cyclophorus affinis* | Myanmar | Freshwater mussel |
